# DfE-DB: A systematic database of 3.8 million human decisions across experience-based tasks

**DOI:** 10.64898/2025.12.12.693971

**Authors:** Yujia Yang, Mikhail S. Spektor, Anna I. Thoma, Ralph Hertwig, Dirk U. Wulff

## Abstract

Learning from experience is central to human decision making, but progress is limited by heterogeneous task designs and fragmented data formats that hinder systematic comparison across studies. We introduce the Decisions From Experience Database (DfE-DB), a structured, open-access resource comprising 3.8 million trial-level decisions from 11,921 participants across 168 studies and 13 paradigms. By harmonizing raw behavioral data and classifying studies along 13 key design features, the DfE-DB enables quantitative comparisons previously obscured by heterogeneous task and data structures. Using the database, we show that choice tendencies—toward higher risk, expected value, or experienced mean—vary substantially across paradigms and are strongly shaped by core design features such as feed-back type, outcome structure, stationarity, and sampling. These features explain substantial cross-study variability and reveal underexplored paradigm variants. By providing a unified and extensible data infrastructure, the DfE-DB facilitates reproducible research on testing the generality of behavioral phenomena and computational models, fostering a more integrated science of decisions from experience.

## Introduction

Understanding how people make decisions under uncertainty is crucial for both research and policy. To study this behavior, the decision sciences often rely on monetary lotteries or other tasks with explicit descriptions of possible outcomes and associated probabilities (Allais, 1953; Savage, 2012; Tversky & Kahneman, 1992). For instance, participants might choose between either €3 guaranteed or €4 with 80% probability and otherwise nothing. However, this approach misses key features of real-world decision making. People are rarely aware of all possible outcomes, let alone their probabilities; instead, they must learn about the options and their consequences through experience. This involves learning from feedback and inferring the generative processes underlying that experience (Spektor & Wulff, 2024). Researchers have developed various laboratory paradigms to study situations that mimic these real-world properties, which we refer to collectively as *decisions from experience* (DfE; Hertwig & Erev, 2009; Hertwig et al., 2004; Wulff et al., 2018). In these paradigms, participants repeatedly choose between options, observe the outcomes, and use that information to update their impressions of the available options. A rapidly growing literature across psychology, economics, and neuroscience (Thoma, Bolenz, et al., 2025) deploys these paradigms under various labels, including *sampling paradigm, probability learning, repeated choices*, and *reinforcement learning* (Dayan & Balleine, 2002; Gonzalez & Mehlhorn, 2016; Palminteri & Lebreton, 2022; Thoma, Newell, & Schulze, 2025). Interest in these paradigms has grown in recent decades, partly due to the discovery of systematic differences between decisions based on description and those based on experience, known as the *description– experience gap* (Hertwig et al., 2004; Wulff et al., 2018). However, this literature is heavily fragmented: Paradigms vary along dimensions that are not systematically characterized, and there is no solid basis for empirically comparing behavior across dimensions. We present the Decisions from Experience Database (DfE-DB), which systematizes decisions-from-experience paradigms, providing structured and open-access data for 3.8 million decisions from 11,921 participants across 168 studies and 13 distinct paradigms. By documenting the diversity of paradigms with respect to design features linked to distinct psychological mechanisms, the DfE-DB provides a platform for integrative research.

Scientific fragmentation is a critical obstacle to cumulative progress in the study of human decision making (Anvari et al., 2025; Núñez et al., 2019; Wulff & Mata, 2025b), preventing collaboration and theoretical integration. Fragmentation can obscure psychologically important effects of seemingly auxiliary design features in behavioral paradigms, such as whether feedback is free or costly, whether options vary in outcomes or only probabilities, and whether options are stationary or dynamically changing (Eckstein et al., 2022; Edwards, 1962). While design features often receive little attention, they shape cognitive processes in distinct ways. For instance, free versus costly feedback determines whether a person faces an exploration–exploitation trade-off, thus separating two distinct classes of paradigms that are typically referred to as *sampling* and *repeated choice* (Hills et al., 2015; Spektor & Wulff, 2024). Documenting these features comprehensively and helping to understand how they influence behavior is the primary goal of the DfE-DB.

The DfE-DB combines datasets from two systematic literature searches. The first search, conducted by Wulff et al. (2018), led to a database containing 1.22 million decisions from sampling paradigms. The second search resulted in a database comprising 2.55 million decisions involving repeated-choice paradigms. See the ‘Method’ section and Supplementary Materials for details on the literature search. Together, these databases represent a broad spectrum of research on decisions from experience.

The DfE-DB provides data in a structured and openly accessible format, presents an initial categorization of paradigms and features, and facilitates reproducible research and systematic comparisons across disciplinary and paradigmatic boundaries. By harmonizing heterogeneous datasets, the database enables new insights into the role of design features(Almaatouq et al., 2024), tests of the generalizability of behavioral patterns, and the development of computational models that account for behavior across paradigms(Binz et al., 2025). Thereby, DfE-DB facilitates a more robust understanding of human decision making that extends beyond individual paradigms. Here, we provide details on the database composition and demonstrate its utility by analyzing the effect of design features and paradigm classes on decision making.

## Method

### Literature Search and Study Identification

The DfE-DB combines the results of two systematic literature searches across two classes of paradigms: sampling and repeated choice. The first search, conducted by Wulff et al. (2018) in December 2015, combined keyword, forward, and backward searches to identify studies reporting data from sampling paradigms. In this paradigm class, participants explore options without cost before making a single consequential decision. The search yielded a total of 29 candidate articles. Additional details on the literature search and data acquisition are presented in Wulff et al. (2018).

We conducted the second search using the same approach to identify studies using repeated-choice paradigms. Unlike sampling paradigms, repeated-choice paradigms are immediately consequential. In other words, choices result directly in monetary gains or losses or other non-monetary outcomes. We selected four seed articles for a backward search and 12 for a forward search (see Table S1). Additionally, we implemented a query using five keywords: ‘repeated choice*’ OR ‘repeated-choice paradigm’ OR ‘feedback paradigm*’ OR ‘forgone outcome*’ OR ‘bandit task*’. Conducted on April 15, 2016, on Web of Science, this search yielded 1,697 articles with full-text availability (see Table S1 for a break-down). We screened each article based on two eligibility criteria: inclusion of (1) raw data from human participants, and (2) repeated choices between lottery-like options. We formulated the latter criterion broadly to include studies involving, for instance, repeated choices between food items. Note that our criteria excluded studies that used the Iowa Gambling Task, as this task is typically deterministic rather than lottery-like. The screening resulted in 155 candidate articles.

### Data Acquisition

We contacted the corresponding authors of all eligible articles to request access to raw trial-level behavioral data and any additional datasets that might meet our inclusion criteria. Of the 29 candidate articles in sampling paradigms and 155 candidate articles in repeated-choice paradigms, we obtained data from 27 and 112 articles, respectively. Three of the articles contained data from both sampling and repeated-choice paradigms, resulting in a total of 136 articles. Upon reviewing the data, we excluded 12 articles due to a lack of trial-by-trial choice data. The data of the remaining 124 articles were subsequently processed using a standardized pipeline. The datasets varied substantially in format (e.g., comma-separated values text files, MATLAB structures, R data, or SPSS structures) and level of documentation. In cases where definitions of variables or task structures were unclear, we followed up with authors to ensure accurate interpretation and reconstruction of the data.

### Data Processing and Standardization

The varying software, structures, and organization of experimental data present major obstacles to data reuse and require reformatting (e.g., choices were stored in a wide format), combining (e.g., each participants’ data were stored in separate files), or flattening (e.g., most MATLAB structures’ trial-level information was organized hierarchically). A key contribution of the DfE-DB is its data format, which standardizes the data representation while accounting for the variability in paradigms and design features.

Data processing involved converting all data into study-level comma-separated value files in a long format, where each row corresponds to a single trial by a single participant and columns encode standardized variables, including study and participant identifiers, problem or option pair identifiers, choice, outcome, response time, and available demographic information. The data processing code is available at (https://github.com/dwulff/DfE-DB).

During processing, we also standardized key variables across studies to ensure semantic consistency. Choice options were encoded alphabetically (e.g., A, B, C), with each letter corresponding to a specific option in the original task; this mapping was held constant across all participants within a study. Outcome variables were harmonized according to their format: numerical outcomes were retained, binary outcomes were recoded as 0/1, and categorical feedback (e.g., food items) was recorded using original labels. The mapping between options and their associated outcomes was encoded in additional study-specific option tables.

### Paradigms and features definition

Research on decisions from experience lacks a systematic overview and a common vocabulary for the many paradigms and their design features. In creating DfE-DB, we sought to provide a first taxonomy. Specifically, we identified 17 distinct paradigm (Table 1) and 13 design feature(Table 2). We further categorized paradigms into paradigm classes by grouping together paradigms that occurred at least three times and showed substantial correspondence. Five paradigms that occurred fewer than three times in our database were classified as ‘other’, implying 13 distinct paradigm classes for our analysis (including the ‘other’ category). The design features were identified based on their use as factors in experimental investigations and on their likely impact on cognition and behavior, as indicated by past research (see ‘How Paradigms and Features Impact Choice’ further below).

**Table 1.** Descriptions of 13 paradigms. Descriptions characterize paradigms relative to the standard repeated-choice paradigm, lottery bandits.

| Paradigm | Explanation |
| --- | --- |
| Lottery bandits | Participants repeatedly choose between options with a small number of possible monetary outcomes with fixed probabilities. Every choice affects the final performance or payoff. |
| Continuous bandits | At least one option has a continuous (i.e., non-discrete) distribution of outcomes. |
| Dynamic bandits | The outcome distribution of at least one option changes across the task. |
| Description bandits | The outcome distribution is explicitly described to participants before they make decisions. |
| Probability learning | Each of two or more options consists of a single outcome, materializing with distinct probabilities. |
| Dynamic probability learning | A probability learning task where the probabilities change across trials. |
| Two-step dynamic probability learning | A probability learning task with a two-step process, where the choice in the first step determines the options available in the second step. |
| Binary prediction | Either one option with two potential outcomes or two options with perfectly dependent outcomes. The task is to predict which outcome will appear or which option will have the target outcome in the current trial. |
| Social binary prediction | A binary prediction task with social context, usually involving two participants. Participants' outcomes are affected by the choices of the participants. |
| Observe-or-bet | Participants can choose to either observe, leading to a non-consequential choice with feedback (i.e., sampling), or bet, leading to a consequential choice without feedback. |
| Free sampling | Participants can sample freely from options to collect information without influencing their performance or payoff. After gathering enough information, participants make one final consequential decision. |
| Regulated sampling | Similar to free sampling, except that after a predetermined number of samples (known or unknown to the participants), participants make one final consequential decision. |
| Other | Other paradigms that do not form a larger paradigm class. Probability learning from observation, probabilistic instrumental learning, correlation detection, exploration task, and probabilistic feedback binary prediction occurred fewer than three times in our database. |

**Table 2.**
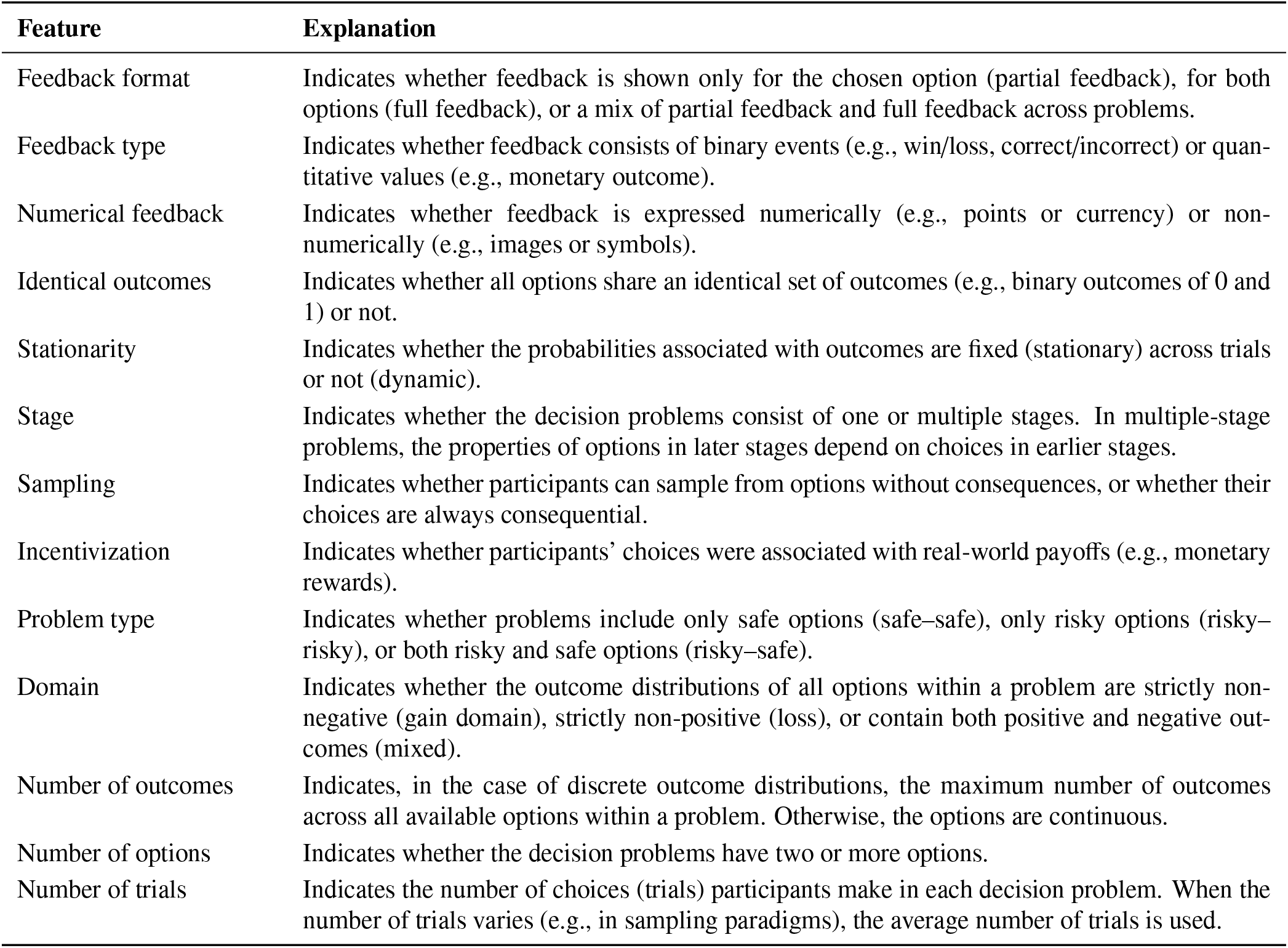
Feature definition. Definitions of key features used to distinguish paradigms. Further details are presented in Table S2

| Feature | Explanation |
| --- | --- |
| Feedback format | Indicates whether feedback is shown only for the chosen option (partial feedback), for both options (full feedback), or a mix of partial feedback and full feedback across problems. |
| Feedback type | Indicates whether feedback consists of binary events (e.g., win/loss, correct/incorrect) or quantitative values (e.g., monetary outcome). |
| Numerical feedback | Indicates whether feedback is expressed numerically (e.g., points or currency) or non-numerically (e.g., images or symbols). |
| Identical outcomes | Indicates whether all options share an identical set of outcomes (e.g., binary outcomes of 0 and 1) or not. |
| Stationarity | Indicates whether the probabilities associated with outcomes are fixed (stationary) across trials or not (dynamic). |
| Stage | Indicates whether the decision problems consist of one or multiple stages. In multiple-stage problems, the properties of options in later stages depend on choices in earlier stages. |
| Sampling | Indicates whether participants can sample from options without consequences, or whether their choices are always consequential. |
| Incentivization | Indicates whether participants' choices were associated with real-world payoffs (e.g., monetary rewards). |
| Problem type | Indicates whether problems include only safe options (safe–safe), only risky options (risky–risky), or both risky and safe options (risky–safe). |
| Domain | Indicates whether the outcome distributions of all options within a problem are strictly non-negative (gain domain), strictly non-positive (loss), or contain both positive and negative outcomes (mixed). |
| Number of outcomes | Indicates, in the case of discrete outcome distributions, the maximum number of outcomes across all available options within a problem. Otherwise, the options are continuous. |
| Number of options | Indicates whether the decision problems have two or more options. |
| Number of trials | Indicates the number of choices (trials) participants make in each decision problem. When the number of trials varies (e.g., in sampling paradigms), the average number of trials is used. |

### Data Sharing

After constructing the database, we contacted all authors who contributed repeated-choice data to seek permission to share their data publicly. The authors of ten articles either declined or did not respond to multiple requests, leaving 90 articles with permission for publication. Note that the authors who contributed data from sampling paradigms previously granted permission for publication in Wulff et al. (2018).

### Data Records

The final DfE-DB comprises data from 114 articles, covering 168 studies and 11,921 participants, with approximately 3.77 million trial-level decisions, with sampling paradigms contributing 27 articles, 40 studies, 4,188 participants, and 1.22 million choices, and repeated-choice paradigms contributing 90 articles, 128 studies, 7,733 participants, and 2.55 million choices; three articles contain both sampling and repeated-choice paradigms. As of 29^th^ January 2026, these articles have been cited a total of 13,220 times according to Semantic Scholar, averaging 8.0 citations per article per year since publication. See Table S3 for an overview of the included articles.

In the final database, each study is documented in two comma-separated-value files: a data table and an options table. The database repository also contains a study-level feature table that documents key study features and other technical details (https://github.com/dwulff/DfE-DB).

## Results

Here, we conduct two sets of exemplary analyses. First, we provide an overview of paradigms and design features and analyze how they interrelate. Second, we analyze how paradigms and design features impact choice with respect to three core preference indicators, and how the effects of paradigms can be accounted for by design features. These analyses highlight how the heterogeneity of behavioral paradigms and design features shape decisions from experience.

### Overview of the Database

Figure 1 illustrates the distribution of studies across paradigms. The paradigms most frequently represented in the database are lottery bandits and free sampling (32 studies each), followed by probability learning (25 studies; see Table 1 for descriptions of paradigms). Although most data come from standard behavioral studies, 23 studies involve psychophysiological measurements (e.g., EEG or fMRI) or special populations.

**Figure 1.**
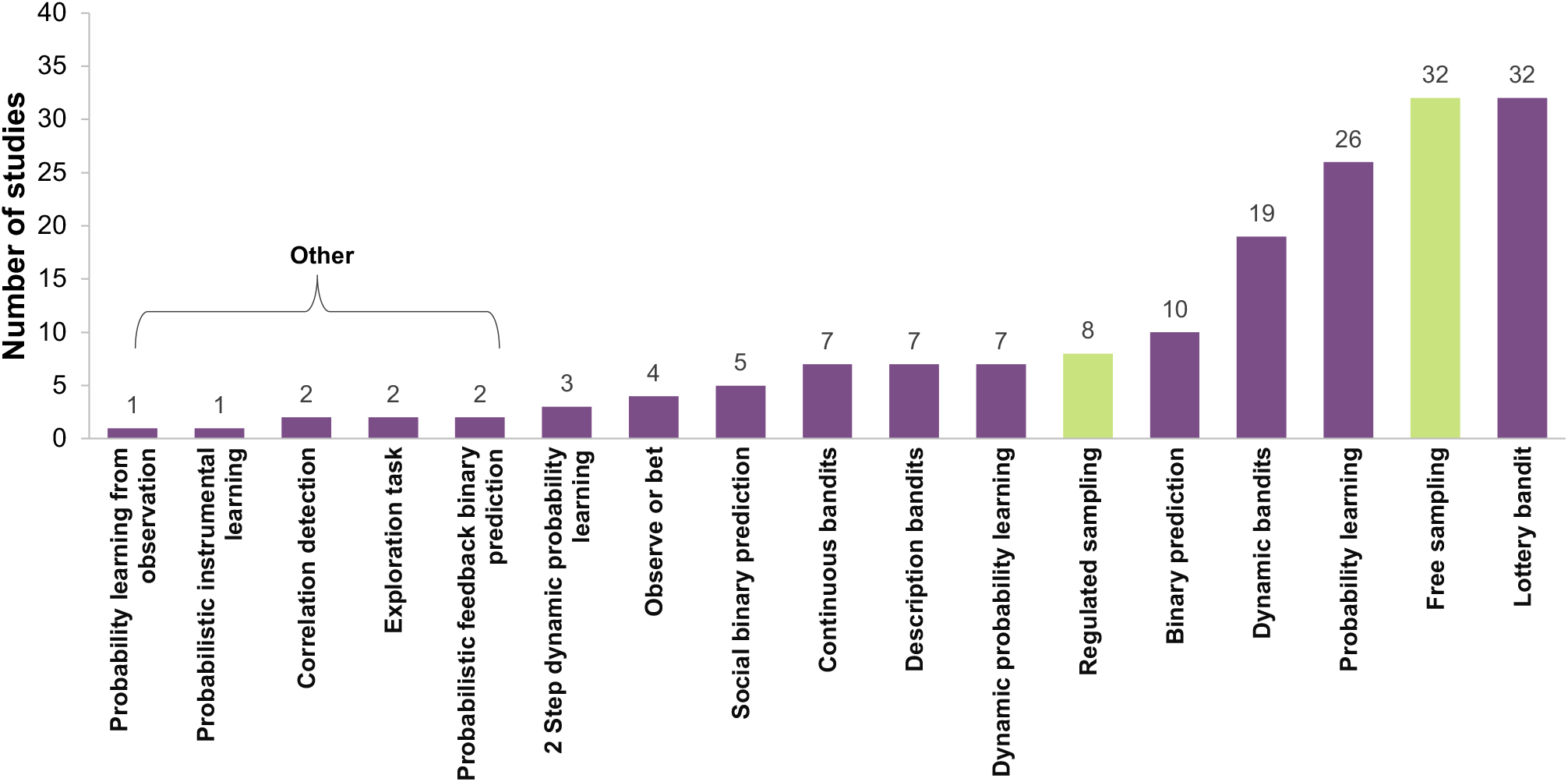
Paradigm frequency. Number of studies included in the DfE-DB for each paradigm class. Tasks with fewer than three studies were grouped together under the classification ‘Other.’ Color indicates paradigm classification: green for sampling paradigms, purple for repeated-choice paradigms. See Table 1 for detailed descriptions of the paradigms.

Figure 2 provides a detailed overview of the 168 studies based on the 13 key design features: feedback format, feedback type, numerical features, identical outcome, stationarity, stage, sampling, incentivization, problem type, domain, number of outcomes, number of options, and number of trials (see Table 2 for descriptions). Although the distribution of these features is skewed and complex, the majority of studies have a single stage (95.24%), involve two-option problems (92.86%) with a maximum of two outcomes per option (86.31%), and present partial (69.64%) and numerical feedback (70.24%). This specific combination characterizes many lottery-bandit, binary-prediction, and probability-learning studies, accounting for 38.10% of the studies in the database. Beyond these common feature combinations, however, feature correlations become more complex.

**Figure 2.**
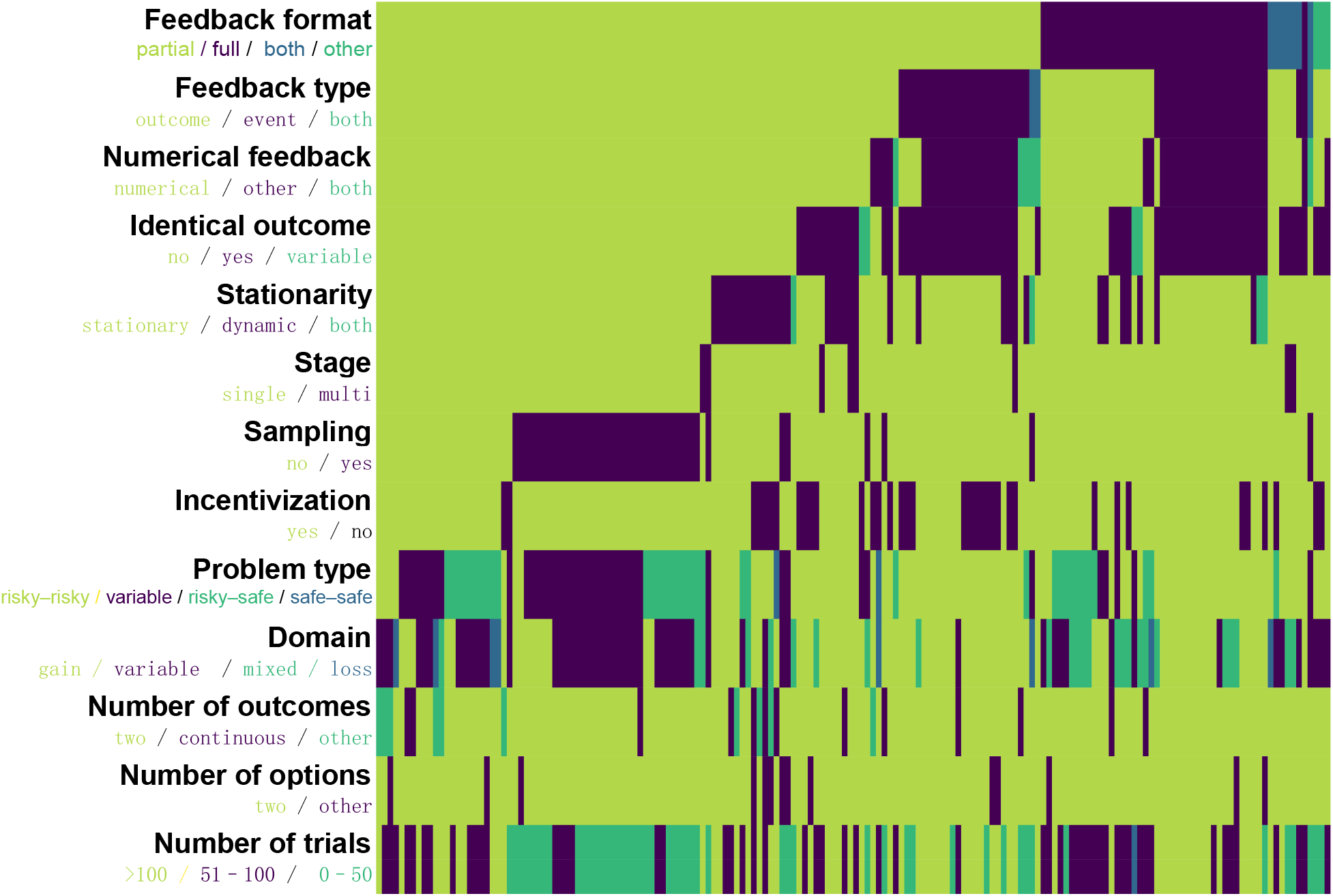
Distribution of design features. Each column refers to one study, with a total of 168 studies.

**Figure 3.**
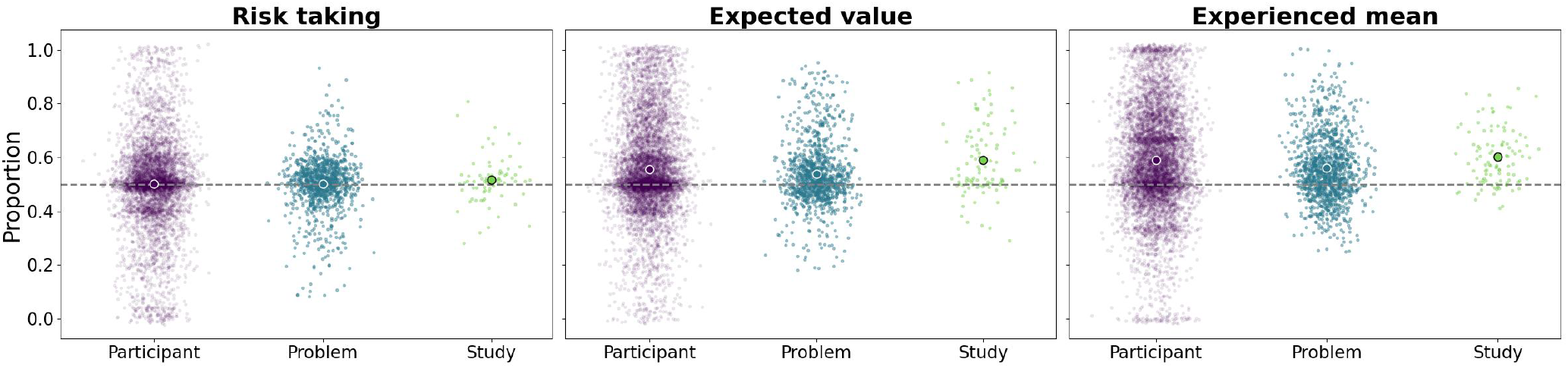
Choice distribution across participants, problems, and studies. Mean and distribution of choice proportions within each level (participant, problem, study). Larger circles indicate the mean of all data points per level.

To identify additional patterns among design features, we calculated multiclass correlations (Cramér’s *V*) for pairs of features. First, problem type was strongly associated with identical outcomes (*V* = .54) and free sampling (*V* =.60). This association is driven by probability-learning paradigms (typically involving identical outcomes) and sampling paradigms (frequently relying on risky–risky problems). Second, the number of trials is strongly associated with sampling (*V* = .68): Participants in sampling paradigms typically freely decide to terminate information sampling and do so, on average, after only 14.1% of the number of trials in non-sampling paradigms. Third, sampling is associated with domain (*V* = .41), as sampling studies are less likely than other types of studies to use decision problems exclusively in the gain domain.

These correlations (see Fig. S1) are critical for two reasons. First, they reveal a lack of orthogonal variation in design features, which can introduce hidden confounding factors that hinder the integration of research across paradigms (Almaatouq et al., 2024). Features such as problem type, sample size, and domain have been found to strongly influence choices in decisions from experience (Hau et al., 2008; Wulff et al., 2018). If not explicitly considered, these features could thus inadvertently affect the cognition and behavior investigated by a study. Second, they imply the existence of feature combinations that are underexplored, pointing to unexamined behavioral patterns.

### How Paradigms and Features Impact Choice

Here, we present an analysis of how paradigms and features affect three core preference indicators: choosing the option with higher variance (reflecting risk-taking attitudes), higher expected value, or higher experienced mean. Risk is a fundamental dimension of decision making, typically involving a trade-off between a larger, less likely outcome and a smaller, more likely one (Hertwig et al., 2019; Mata et al., 2018; Wulff & Mata, 2022). Expected value often serves as a normative benchmark for assessing learning or decision-making performance in the context of risky choice (He et al., 2022; Palminteri, 2025; Rolls et al., 2008). Experienced mean is a common empirical benchmark—often referred to as the natural mean heuristic—that has been found to explain a substantial proportion of choices, frequently exceeding the explanatory power of expected-value maximization (Hertwig, 2015; Wulff et al., 2018).

We evaluated these three indicators for each decision problem—that is, for each unique constellation of outcomes and probabilities across the decision options. We aimed to address three questions: Are there consistent choices across the three indicators? How do choices vary as a function of paradigm? And finally, can design features explain the potential variability in choices? For simplicity, we limited subsequent analyses to cases with no more than two options and no more than two outcomes per option (covering 58.18% of all choices in the database).

### Consistency of Choices in Decisions From Experience

Our analysis of whether there are systematic tendencies across participants, problems, and studies across the three indicators establishes a baseline for the upcoming analyses and identifies the primary sources of variability in choice. On average, participants in decisions from experience exhibit neutrality toward risk (50%) and moderate tendencies to choose higher expected-value (53%) and experienced-mean (55%) options. Choices vary significantly at all three levels, though to vastly different extents. The participant level shows the largest variance, ranging, for each indicator, from a strict preference in favor to a strict preference against. However, after accounting for the variance-reducing effects of aggregation (via the central limit theorem), higher levels of aggregation show greater underlying variation. We estimated population variances of 0.30 for participants (*N* = 7, 298), 0.65 for problems (*N* = 1, 210), and 1.39 for studies (*N* = 104). Note that the variances exceed 1, as the data violate the assumption of a single shared population. These estimates suggest that study design systematically influences choice.

### The Role of Paradigms and Study Features

What accounts for the preference variability observed across problems and studies? Figure 4 displays average choice proportions by decision problem and between paradigms, revealing a pattern of choice as a function of paradigms that is highly similar between the three indicators. Sampling and bandit paradigms (excluding dynamic bandits) show minimal deviation from the 50% baseline. In contrast, probability-learning, binary-prediction, and dynamic-bandit paradigms exhibit medium-to-large average choice proportions favoring higher risk, higher expected value, and higher experienced mean (note that risk is irrelevant in binary-prediction paradigms, as the options in these tasks carry identical risk). However, the pattern is not identical across indicators. For instance, choice proportions for lottery bandits deviate from chance level only for higher experienced-mean choices, not for risk or expected value.

**Figure 4.**
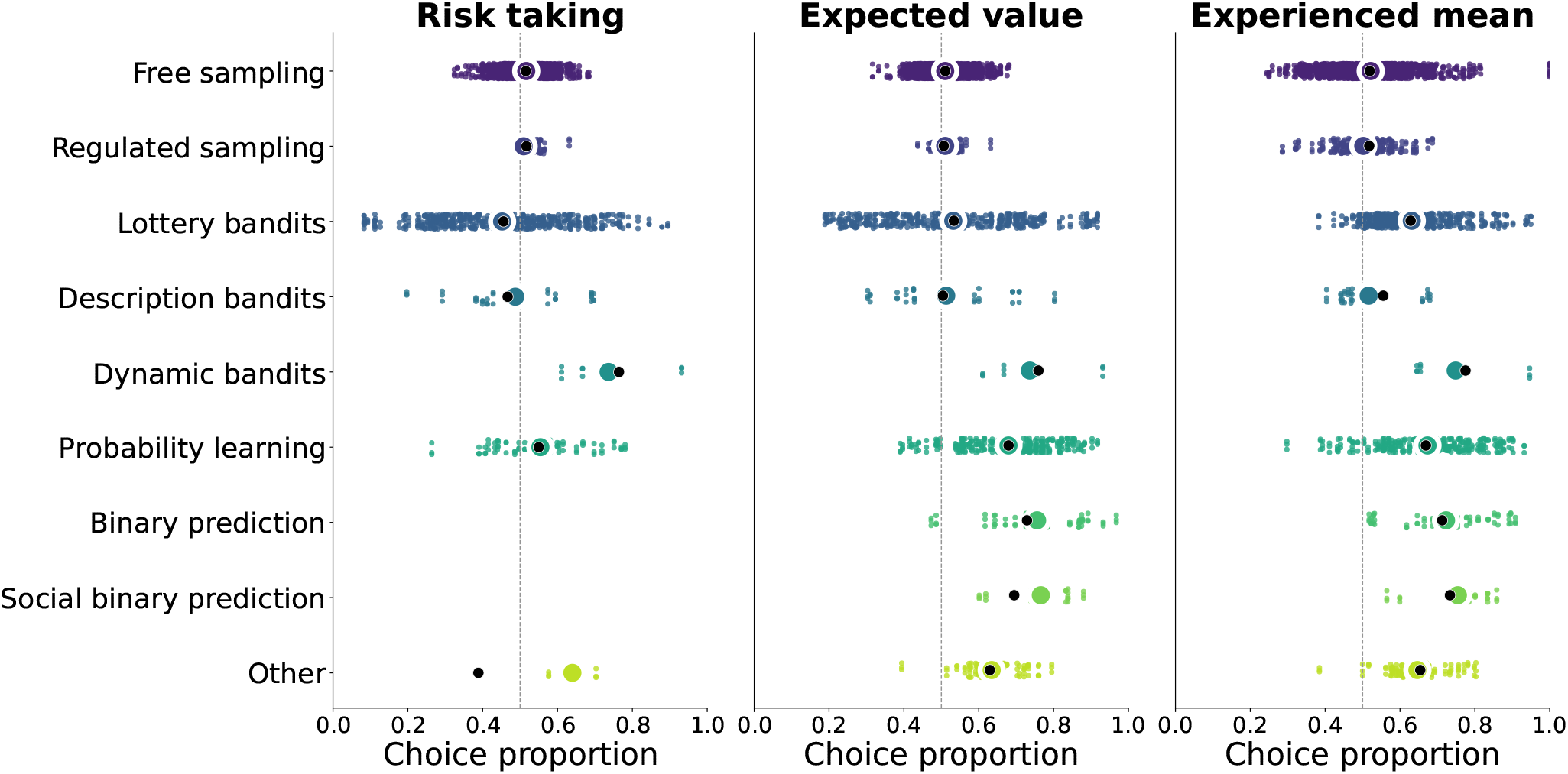
Choice proportion distribution for different paradigms. Colored circles indicate the mean value of the experimental data; black dots indicate the mean predicted values from the feature random forest model.

Figure 5 illustrates choice tendencies as a function of nine of the 13 design features. Three features—stage, number of outcomes, and number of options—were constant across the subset of data analyzed here and thus excluded. Furthermore, feedback type and numerical feedback were largely equivalent, and thus, only feedback type is analyzed here. The results show that feedback type and identical outcome had particularly strong effects, which dictate the structure of decision problems: When feedback is identical across options, options differ solely in terms of probability.

**Figure 5.**
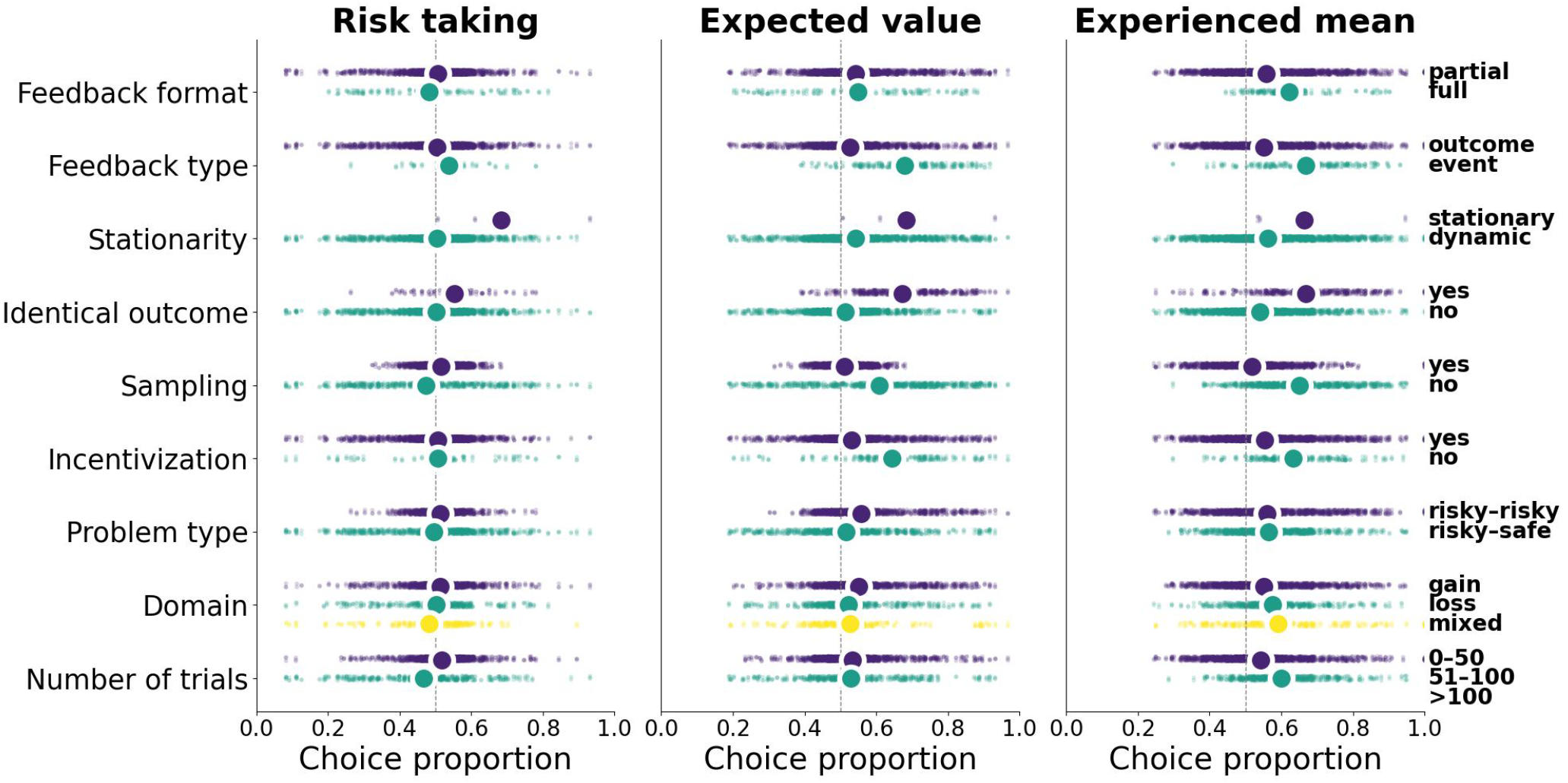
Choice proportion distribution for key design features. Colored circles indicate the mean value of the experimental data.

This results in one option dominating the other unless the probabilities are equal, which may explain people’s stronger tendencies towards choosing options with higher risk, expected value, and experienced mean in probability-learning and binary-prediction paradigms.

Stationarity and sampling also exhibit notable effects. Stationarity relates to the stability of option properties over time. In the absence of stationarity (i.e., in dynamic environments), options can become more easily distinguishable as the problem characteristics shift and one option—usually the riskier option—emerges as clearly superior in terms of expected value and experienced mean. This increased distinguishability likely drives the stronger preferences seen in dynamic bandits. Sampling refers to whether participants can explore options without consequences. When each choice yields a gain or loss, participants are incentivized to select options that offer the higher average rewards according to their experience. This can account for the systematic choice tendency in favor of the higher experienced mean in lottery bandits, where—unlike description bandits—only experienced information is available. In contrast, when there are no consequences to sampling, there is less incentive to maximize the experienced mean during sampling, which explains the baseline results in standard sampling paradigms.

We also examined whether other features contributed to explaining the choice patterns displayed in Figures 4 and 5 by running two random forest models for each preference indicator: one that used only the four features discussed above (feedback type, identical outcome, stationarity, and sampling) and another that used all nine non-redundant features. The four features accounted for 8.0% (risk taking), 11.4% (expected value), and 21.2% (experienced mean) of the variance across decision problems. The nine-feature model accounted for 5.8% (risk taking), 23.1% (expected value), and 34.3% (experienced mean) of the variance, indicating that the additional features made important contributions to the variance in expected mean and experienced mean. In general, design features capture considerable variability in choice proportions with respect to risk, expected value, and experienced mean. Notably, as shown in Figure 4, the nine features together account for almost all of the variation in average choice proportions across paradigms.

Collectively, these results confirm substantial differences in people’s decision making across the paradigms in our database. These differences can largely be attributed to design features that past work has shown to correspond to distinct psychological processes. Overall, this analysis demonstrates the usefulness of the DfE-DB in illuminating how paradigms and design features affect experience-based choice.

## Discussion

The study of how people make decisions from experience is a large and influential field (Hertwig & Erev, 2009; Olschewski et al., 2024; Plonsky et al., 2025; Wulff et al., 2018) whose progress has been slowed by fragmentation (Thoma, Bolenz, et al., 2025). Different subfields have often worked in parallel, relying on diverse paradigms under various labels, impeding theoretical integration and the generalizability of findings. DfE-DB responds to this challenge. By standardizing 3.8 million decisions across 13 paradigms, the database provides the foundational infrastructure needed to unify this fragmented landscape.

A key obstacle to cumulative progress in decision-making research is that theories are often not tested in other paradigms than the one that led to their development(Almaatouq et al., 2024; Binz et al., 2025; Wulff & Mata, 2025a; Yarkoni, 2022). Using DfE-DB, researchers can now rigorously test the generalizability of documented behavioral patterns and computational models. Answering questions such as whether the same reinforcement learning model can equally predict behavior in classic lottery-bandit, probability-learning, and dynamic-bandit tasks is essential for more robust and comprehensive theories of human learning and choice.

Our exemplary analysis revealed correlations between design features across paradigms that may account for discrepancies in the literature: Seemingly minor design features—such as feedback format or whether outcomes are constant across options—can systematically co-vary with paradigms and profoundly influence choice. In this regard, our results confirm past evidence linking design features and decision making. For instance, feedback format has been linked to distinct neural activations (Lohrenz et al., 2007) and risk taking (Camilleri & Newell, 2011b; Shiravand et al., 2025; Yechiam & Busemeyer, 2006), problem type has been linked to information search strategies (Spektor & Wulff, 2024) and the description–experience gap (Hertwig & Erev, 2009; Wulff et al., 2018), and number of outcomes has been linked to use of distinct memory strategies (Konstantinidis et al., 2022; Olschewski et al., 2024). DfE-DB enables researchers to move beyond treating paradigms as monolithic and explore how individual design features influence cognition. By highlighting feature combinations that are currently underexplored, our database further identifies novel experimental designs that could uncover unobserved behavioral phenomena.

Some limitations warrant mention. First, the database mostly reflects published studies, which may overrepresent significant effects. Note, however, that studies have not been selected concerning any particular hypothesis. Second, our list of 13 features, while comprehensive, is not exhaustive; features such as time pressure or instruction framing may also shape cognition and behavior. We did not include these features because they are either less frequently implemented (time pressure) or less frequently reported (framing). Third, the correlational nature of cross-study comparisons limits causal inference—targeted experiments that manipulate in-dividual features are essential. Despite these limitations, the database provides a broad empirical foundation that can inform experimental follow-up studies.

The release of the DfE-DB is a starting point for systematizing the analysis of decisions from experience. We encourage the community to use this resource not only for secondary analysis but also as a foundation for future open science and sharing initiatives. By adopting the proposed data format and feature classification, researchers can ensure that their new data can be easily integrated into DfEDB, transforming it into a living, evolving resource. We invite researchers to share new data via the project’s GitHub repository (https://github.com/dwulff/DfE-DB). Such a collaborative effort can accelerate discovery and enhance the replicability and transparency of research on decisions from experience by providing an empirical foundation for comparing and integrating findings. We hope DfE-DB will help the field move toward a robust and integrated science of decisions from experience.

## Supporting information

all_sipplementary materials

## Acknowledgments

We thank the authors of the original research for sharing their data and Deb Ain for editing the manuscript.

## Author contributions

Conceptualization: M.S.S., R.H., D.U.W.; Validation: Y.Y., M.S.S., D.U.W.; Formal analysis: Y.Y.; Data Curation: Y.Y., M.S.S., A.I.T., D.U.W.; Database organization: Y.Y., M.S.S., D.U.W.; Visualization: Y.Y., D.U.W.; Supervision: D.U.W.; Writing - Original Draft: Y.Y., D.U.W.; Writing - Review & Editing: Y.Y., A.I.T., M.S.S., R.H., D.U.W.; Funding acquisition: R.H. & D.U.W.

## Data and materials availability

The database is stored in a GitHub repository (https://github.com/dwulff/DfE-DB). The data is currently made available to data providers and will be shared publicly upon article acceptance. Code reproducing all analyses in this paper is available at https://github.com/Yujia-Y/DfE-DB-Analysis.

