## Supplementary material for "DfE-DB: A systematic database of 3.8 million human decisions across experience-based tasks": all_sipplementary materials

#### A Literature Search Strategy: Repeated Choice

Following the strategy Wulff et al. (2018) used for the sampling paradigm, our literature search for the repeated-choice paradigm employed three search strategies: forward citation search, backward citation search, and keyword search. Table S1 provides details on each strategy, including the articles used for the forward and backward search, the search queries for the keyword search, and the number of hits.

Table S1: **Summary of literature search results** across forward citation search, backward citation search, and keyword search.

| Search Type / Item | Hits | New unique | Retrieved |
|---|---|---|---|
| **Forward Citation Search** | | | |
| Bush and Mostellar (1951)(Bush & Mosteller, 1951) | 263 | 263 | 237 |
| Edwards (1961)(Edwards, 1961) | 191 | 191 | 168 |
| Katz (1964)(Katz, 1964) | 16 | 16 | 14 |
| Myers et al. (1965)(J. L. Myers et al., 1965) | 23 | 18 | 16 |
| Rapoport (1964)(Rapoport, 1964) | 15 | 11 | 10 |
| Tversky and Edwards (1966)(Tversky & Edwards, 1966) | 44 | 43 | 37 |
| Busemeyer (1982)(Busemeyer, 1982) | 21 | 18 | 18 |
| Yechiam and Busemeyer (2005)(Yechiam & Busemeyer, 2005) | 82 | 74 | 69 |
| Barron and Erev (2003)(Barron & Erev, 2003) | 226 | 192 | 183 |
| Gonzalez et al. (2003)(Gonzalez et al., 2003) | 107 | 71 | 66 |
| Dayan and Balleine (2002)(Dayan & Balleine, 2002) | 348 | 347 | 329 |
| Niv (2009)(Niv, 2009) | 84 | 81 | 78 |
| **Backward Citation Search** | | | |
| Plonsky et al. (2015)(Plonsky et al., 2015) | 90 | 65 | 55 |
| Gonzalez and Dutt (2011)(Gonzalez & Dutt, 2011) | 67 | 40 | 34 |
| Frank et al. (2015)(Frank et al., 2015) | 47 | 45 | 44 |
| Lee et al. (2012)(D. Lee et al., 2012) | 149 | 140 | 134 |
| **Keyword Search** | | | |
| ‘repeated choice*’ | 167 | 146 | 115 |
| ‘repeated-choice paradigm’ | 3 | 0 | 0 |
| ‘Feedback paradigm’ | 98 | 95 | 84 |
| ‘Forgone outcome*’ | 4 | 1 | 1 |
| ‘bandit task*’ | 50 | 9 | 5 |
| **Summary** | 2095 | 1866 | 1697 |

*Note.* “New Unique” indicates articles not covered by previous search procedures. “Retrieved” denotes articles for which the full text was available.

#### B Features Definition

Table S2: **Description of features.** This table provides definitions of all study and design features characterizing each study in our database. Some feature values were grouped together for the analysis in the main text to increase simplicity. For instance, the feedback formats "none" and "other" were grouped into one group labeled "other".

| Feature | Description |
|---|---|
| **study** | Study identifier. Concatenation of the first letters of the authors' last names, publication year, and study index. Example: AK2011_1 refers to the first study of Avrahami and Kareev (2011). |
| **paradigm** | Name of the paradigm category assigned to the study. |
| **n_participants** | Total number of participants in the experiment. |
| **n_problems** | Total number of problems in the experiment. |
| **n_problems_per_participant** | Number of problems completed by each participant. If different participants completed different numbers of problems, the largest number is shown. |
| **problems_differ** | "yes" if the number of problems per participant differs; otherwise "no". |
| **n_trials** | Number of choices per problem. If the number varies between problems, the mean is returned. |
| **trials_differ** | "yes" if the number of trials per problem differs; otherwise "no". |
| **n_options_per_problem** | Number of options per problem. If the number varies, the largest number is returned. |
| **n_outcomes_per_option** | Number of outcomes per option. If outcomes differ between options, the largest number is returned. "continuous" indicates continuous outcomes. |
| **stationarity** | **stationary**: Outcomes and probabilities are constant across trials or choices. |
| | **dynamic**: Outcomes and/or probabilities change across trials or choices. |
| **identical outcomes** | "yes" if all options share an identical set of outcomes; otherwise "no". |
| **feedback_format** | **partial**: Only the outcome of the chosen option was displayed. |
| | **full**: Outcomes of chosen and non-chosen options were displayed. |
| | **both**: Partial feedback in some conditions and full in others. |
| | **none**: No feedback from either option. |
| | **other**: Other feedback types not included above. |
| **numerical_feedback** | **numerical**: Feedback is a number (numeric value or points). |
| | **graphical**: Feedback is an image (e.g., a picture of a coin). |
| **feedback_type** | **event**: Binary feedback (e.g., 0/1, true/false). |
| | **outcome**: Numerical feedback (whole or real numbers). |
| **problem_type** | **safe_safe**: Only safe options. |
| | **risky_safe**: Mix of safe and risky options. |

Continued on next page

| Feature | Description |
|---|---|
| | **risky_risky**: Only risky options. |
| | **variable**: Varies between problems. |
| **problem_domain** | **gain**: Only positive outcomes. |
| | **loss**: Only negative outcomes. |
| | **mixed**: Contains both gains and losses. |
| | **variable**: Domain differs between problems. |
| **problem_stages** | **single**: Choices lead directly to payoff. |
| | **multi**: Choices lead to higher-order problems or follow-up phases. |
| **sampling** | "yes" if participants can sample without consequences; otherwise "no". |
| **n_conditions** | Number of conditions. |
| **design** | "between" if conditions vary between participants; otherwise "within". |
| **study_context** | **behavioral** |
| | **clinical** |
| | **EEG** |
| | **MRI (incl. fMRI)** |
| | **transcranial** |
| | **pharmacological** |
| | **psychophysiological** |
| | **genetics** |
| **decisions_from_description** | "yes" if the study also involved description-based choices. |
| **incentivization** | "yes" if choices were incentivized; otherwise "no". |

### C Features Relationships

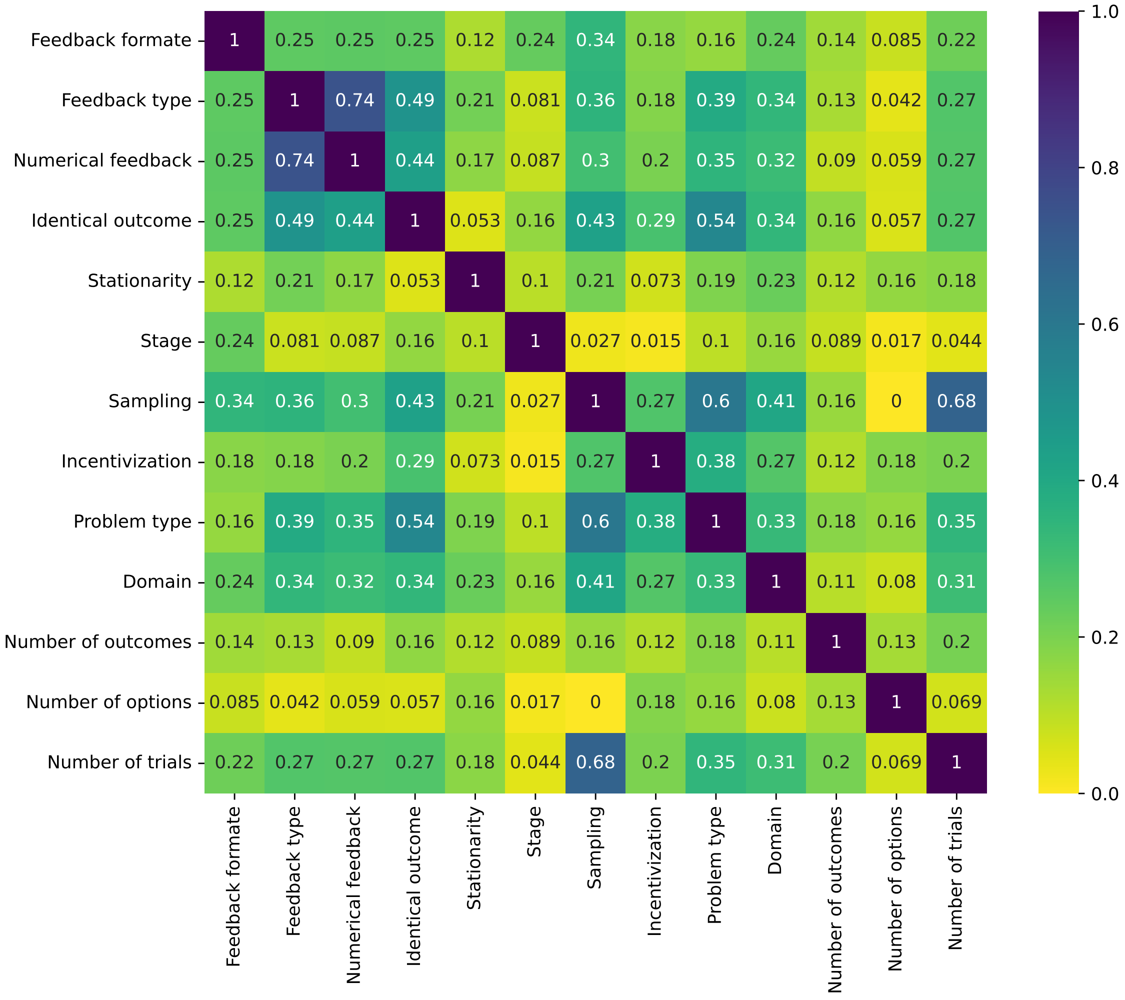


Figure S1: **Multi-Class Correlation Heatmap for All Features.** This figure shows the multi-class correlations (Cramer's $V$) between design features.

### D Papers Included in the Database

Table S3: **Paper included in the database.** Include the author, the short name we used in database, the number of participants, problems, total trials, and studies included in this paper.

| Article | Label | $N$ participants | $N$ problems | $N$ trials | $N$ studies |
|---|---|---|---|---|---|
| Avrahami, J., & Kareev, Y. (2011)(Avrahami & Kareev, 2011) | AK2011 | 95 | 2 | 9500 | 2 |
| Avrahami, J. et al. (2014)(Avrahami et al., 2014) | AKH2014 | 60 | 2 | 4784 | 1 |
| Ashby, N. J., & Rakow, T. (2016)(Ashby & Rakow, 2016) | AR2016 | 51 | 14 | 28560 | 1 |

Continued on next page

| Article | Label | $N$ participants | $N$ problems | $N$ trials | $N$ studies |
|---|---|---|---|---|---|
| Bereby-Meyer, Y., & Erev, I. (1998)(Bereby-Meyer & Erev, 1998) | BE1998 | 42 | 3 | 21000 | 1 |
| Barron, G., & Erev, I. (2003)(Barron & Erev, 2003) | BE2003 | 240 | 12 | 97600 | 5 |
| Biele, G.et al. (2009)(Biele et al., 2009) | BEE2009 | 94 | 22 | 70400 | 2 |
| Beevers, C. G. et al. (2013)(Beevers et al., 2013) | BWGN2013 | 95 | 2 | 15200 | 1 |
| Barron, G., & Yechiam, E. (2009)(Barron & Yechiam, 2009) | BY2009 | 40 | 2 | 16000 | 1 |
| Cooper, J. C. et al. (2012)(J. C. Cooper et al., 2012) | CDFO2012 | 16 | 10 | 6332 | 1 |
| Cooper, J. A. et al. (2014).(J. A. Cooper et al., 2014) | CGDW2014 | 75 | 1 | 11250 | 1 |
| Camilleri, A. R., & Newell, B. R. (2009) (Camilleri & Newell, 2009) | CN2009a | 80 | 8 | 8696 | 1 |
| Camilleri, A. R., & Newell, B. (2009) (Camilleri & Newell, 2009) | CN2009b | 40 | 10 | 4770 | 1 |
| Camilleri, A. R., & Newell, B. R. (2011) (Camilleri & Newell, 2011a) | CN2011a | 102 | 20 | 14998 | 2 |
| Camilleri, A. R., & Newell, B. R. (2011)(Camilleri & Newell, 2011b) | CN2011b | 80 | 8 | 32000 | 2 |
| Camilleri, A. R., & Newell, B. R. (2013) (Camilleri & Newell, 2013) | CN2013 | 203 | 32 | 64960 | 1 |
| Doll, B. B. et al. (2016) (Doll et al., 2016) | DBDF2016 | 171 | 2 | 50643 | 1 |
| Dombrovski, A. Y. et al. (2015) (Dombrovski et al., 2015) | DSCA2015 | 47 | 1 | 14121 | 1 |
| Erev, I. et al. (2010) (Erev et al., 2010) | EERH2010 | 100 | 60 | 120000 | 1 |
| Erev, I. et al. (2008) (Erev et al., 2008) | EEY2008 | 145 | 12 | 57820 | 2 |
| Fleischhut, N. et al. (2014) (Fleischhut et al., 2022) | FAOV2014 | 89 | 12 | 36098 | 1 |
| Frey, R. et al. (2014) (Frey et al., 2014) | FHR2014 | 161 | 13 | 47987 | 2 |
| Fatás, E. et al. (2011) (Fatás et al., 2011) | FJM2011 | 64 | 1 | 9216 | 1 |
| Frey, R. et al. (2015) (Frey et al., 2015) | FMH2015 | 191 | 96 | 202042 | 2 |
| Frank, M. J. et al. (2007). (Frank, Moustafa, et al., 2007) | FMHC2007 | 42 | 2 | 6368 | 1 |

Continued on next page

| Article | Label | $N$ participants | $N$ problems | $N$ trials | $N$ studies |
|---|---|---|---|---|---|
| Frank, M. J. et al. (2007) (Frank, Santamaria, et al., 2007) | FSRW2007 | 46 | 1 | 18641 | 1 |
| Gershman, S. J. (2016) (Gershman, 2016) | G2016 | 205 | 12 | 20500 | 2 |
| Glöckner, A. et al. (2012) (Glöckner et al., 2012) | GFHA2012 | 22 | 59 | 42055 | 1 |
| Glöckner, A. et al. (2016) (Glöckner et al., 2016) | GHHF2016 | 114 | 250 | 308400 | 3 |
| Gonzalez, C., & Mehlhorn, K. (2016) (Gonzalez & Mehlhorn, 2016) | GM2016 | 925 | 6 | 42902 | 2 |
| Gershman, S. J., & Niv, Y. (2015) (Gershman & Niv, 2015) | GN2015 | 29 | 15 | 18504 | 2 |
| Gaissmaier, W., & Schooler, L. J. (2008) (Gaissmaier & Schooler, 2008) | GS2008 | 219 | 2 | 126144 | 2 |
| Gaissmaier, W. et al. (2006) (Gaissmaier et al., 2006) | GSR2006 | 160 | 4 | 61440 | 2 |
| Hertwig, R. et al. (2004) (Hertwig et al., 2004) | HBWE2004 | 50 | 6 | 3136 | 1 |
| Hochman, G., & Erev, I. (2013) (Hochman & Erev, 2013) | HE2013 | 73 | 12 | 14597 | 2 |
| Hadar, L., & Fox, C. R. (2009) (Hadar & Fox, 2009) | HF2009 | 111 | 3 | 6660 | 1 |
| Harman, J. L., & Gonzalez, C. (2015) (Harman & Gonzalez, 2015) | HG2015 | 200 | 3 | 26998 | 2 |
| Hills, T. T. et al. (2013) (Hills et al., 2013) | HNG2013 | 64 | 5 | 7049 | 1 |
| Hertwig, R., & Pleskac, T. J. (2010) (Hertwig & Pleskac, 2010) | HP2010 | 88 | 12 | 15769 | 1 |
| Hau, R. et al. (2008) (Hau et al., 2008) | HPH2008 | 123 | 18 | 36376 | 3 |
| Itthipuripat, S. et al. (2015) (Itthipuripat et al., 2015) | ICRS2015 | 54 | 2 | 103300 | 2 |
| Jessup, R. K. et al. (2008) (Jessup et al., 2008) | JBB2008 | 29 | 2 | 3334 | 1 |
| Jessup, R. K. et al. (2010) (Jessup et al., 2010) | JBB2010 | 20 | 4 | 2383 | 1 |
| Jocham, G. et al. (2011) (Jocham et al., 2011) | JKU2011 | 16 | 2 | 11258 | 1 |
| Kool, W. et al. (2016) (Kool et al., 2016) | KCG2016 | 406 | 2 | 48607 | 2 |
| Knox, W. B. et al. (2012) (Knox et al., 2012) | KOSL2012 | 139 | 3 | 21500 | 1 |

Continued on next page

| Article | Label | $N$ participants | $N$ problems | $N$ trials | $N$ studies |
|---|---|---|---|---|---|
| Kellen, D. et al. (2016) (Kellen et al., 2016) | KPH2016 | 104 | 114 | 249506 | 1 |
| Kim, H. et al. (2006) (Kim et al., 2006) | KSO2006 | 16 | 4 | 5440 | 1 |
| Lejarraga, T. (2010) (Lejarraga, 2010) | L2010 | 124 | 7 | 11817 | 1 |
| Li, J. et al. (2011) (J. Li et al., 2011) | LDP2011 | 20 | 1 | 3200 | 1 |
| Lejarraga, T., & Gonzalez, C. (2011) (Lejarraga & Gonzalez, 2011) | LG2011 | 92 | 2 | 18300 | 1 |
| Lejarraga, T. et al. (2014) (Lejarraga et al., 2014) | LLG2014 | 80 | 6 | 24000 | 1 |
| Li, C. T. et al. (2014) (C.-T. Li et al., 2014) | LLLH2014 | 69 | 2 | 33120 | 1 |
| Lefebvre, G. et al. (2017) (Lefebvre et al., 2017) | LLMB2017 | 85 | 2 | 8087 | 2 |
| Lejarraga, T. et al. (2016) (Lejarraga et al., 2016) | LPFH2016 | 30 | 9 | 10451 | 1 |
| Ludvig, E. A., & Spetch, M. L. (2011) (Ludvig & Spetch, 2011) | LS2011 | 61 | 10 | 11712 | 1 |
| Lee, J. C., & Tomblin, J. B. (2012) (J. C. Lee & Tomblin, 2012) | LT2012 | 33 | 2 | 11831 | 1 |
| Mehlhorn, K. et al. (2014) (Mehlhorn et al., 2014) | MBDG2014 | 294 | 16 | 5614 | 1 |
| Morris, R. W. et al. (2014) (Morris et al., 2014) | MDGB2014 | 20 | 6 | 22095 | 1 |
| Munichor, N. et al. (2006) (Munichor et al., 2006) | MEL2006 | 78 | 2 | 4800 | 2 |
| Myers, C. E. et al. (2016) (C. E. Myers et al., 2016) | MSBL2016 | 80 | 4 | 18880 | 1 |
| Moustafa, A. A. et al. (2015) (Moustafa et al., 2015) | MSM2015 | 199 | 4 | 28560 | 1 |
| Niv, Y. et al. (2015) (Niv et al., 2015) | NDGG2015 | 22 | 1 | 17258 | 1 |
| Nevo, I., & Erev, I. (2012) (Nevo & Erev, 2012) | NE2012 | 76 | 14 | 44237 | 2 |
| Niv, Y. et al. (2012) (Niv et al., 2012) | NEDO2012 | 16 | 1 | 3601 | 1 |
| Noguchi, T., & Hills, T. T. (2016) (Noguchi & Hills, 2016) | NH2016 | 232 | 1572 | 22529 | 2 |
| Newell, B. R. et al. (2013) (Newell et al., 2013) | NKJR2013 | 100 | 1 | 15600 | 1 |

Continued on next page

| Article | Label | *N* participants | *N* problems | *N* trials | *N* studies |
|---|---|---|---|---|---|
| Navarro, D. J. et al. (2016) (Navarro et al., 2016) | NNS2016 | 644 | 31 | 169971 | 2 |
| O'Doherty, J. et al. (2004) (O'Doherty et al., 2004) | ODSD2004 | 12 | 1 | 1651 | 1 |
| Otto, A. R. et al. (2014) (Otto et al., 2014) | OKML2014 | 75 | 3 | 15000 | 2 |
| Otto, A. R. et al. (2011) (Otto et al., 2011) | OTM2011 | 160 | 1 | 49623 | 1 |
| Phillips, N. D. et al. (2014) (Phillips et al., 2014) | PHKA2014 | 180 | 20 | 5479 | 1 |
| Palminteri, S. et al. (2016) (Palminteri et al., 2016) | PKCB2016 | 38 | 4 | 2941 | 1 |
| Palminteri, S. et al. (2015) (Palminteri et al., 2015) | PKJC2015 | 28 | 16 | 10752 | 1 |
| Palminteri, S. et al. (2017) (Palminteri et al., 2017) | PLKB2017 | 40 | 2 | 7680 | 2 |
| Rakow, T. et al. (2008) (Rakow et al., 2008) | RDN2008 | 80 | 12 | 7796 | 1 |
| Ratcliff, R., & Frank, M. J. (2012) (Ratcliff & Frank, 2012) | RF2012 | 30 | 1 | 39483 | 1 |
| Rosati, A. G., & Hare, B. (2016) (Rosati & Hare, 2016) | RH2016 | 125 | 4 | 1500 | 2 |
| Rakow, T., & Miler, K. (2009) (Rakow & Miler, 2009) | RM2009 | 92 | 10 | 34720 | 2 |
| Rakow, T. et al. (2015) (Rakow et al., 2015) | RNW2015 | 151 | 12 | 181440 | 2 |
| Rakow, T. et al. (2010) (Rakow et al., 2010) | RNZ2010 | 80 | 12 | 33480 | 2 |
| Rakow, T., & B. Rahim, S. (2010) (Rakow & B. Rahim, 2010) | RR2010 | 324 | 16 | 26400 | 3 |
| Speekenbrink, M., & Konstantinidis, E. (2015) (Speekenbrink & Konstantinidis, 2015) | SK2015 | 80 | 4 | 15800 | 1 |
| Shin, Y. S. et al. (2014) (Shin et al., 2014) | SKH2014 | 34 | 2 | 6406 | 2 |
| Selbing, I. et al. (2014) (Selbing et al., 2014) | SLO2014 | 40 | 1 | 5481 | 1 |
| Schulze, C., & Newell, B. R. (2015) (Schulze & Newell, 2015) | SN2015 | 180 | 3 | 36000 | 3 |
| Sims, C. R. et al. (2013) (Sims et al., 2013) | SNJG2013 | 12 | 1 | 9600 | 1 |
| Silberberg, A. et al. (2013) (Silberberg et al., 2013) | SPAF2013 | 20 | 4 | 2520 | 1 |

Continued on next page

| Article | Label | $N$ participants | $N$ problems | $N$ trials | $N$ studies |
| --- | --- | --- | --- | --- | --- |
| Skvortsova, V. et al. (2014) (Skvortsova et al., 2014) | SPP2014 | 20 | 1 | 5760 | 1 |
| Schurr, A. et al. (2014) (Schurr et al., 2014) | SRE2014 | 47 | 1 | 4700 | 1 |
| Schulze, C. et al. (2015) (Schulze et al., 2015) | SRN2015 | 100 | 2 | 40000 | 2 |
| Stillwell, D. J., & Tunney, R. J. (2009) (Stillwell & Tunney, 2009) | ST2009 | 21 | 1 | 20740 | 1 |
| Teoderescu, K. et al. (2013) (Teoderescu et al., 2013) | TAE2013 | 110 | 12 | 43998 | 3 |
| Teodorescu, K., & Erev, I. (2014) (Teodorescu & Erev, 2014) | TE2014 | 120 | 6 | 12000 | 2 |
| Turi, Z. et al. (2015) (Turi et al., 2015) | TMOP2015 | 16 | 2 | 11422 | 1 |
| Turi, Z. et al. (2017) (Turi et al., 2017) | TMPA2017 | 29 | 2 | 30898 | 1 |
| Ungemach, C. et al. (2009) (Ungemach et al., 2009) | UCS2009 | 247 | 12 | 31260 | 2 |
| Valentin, V. V. et al. (2007) (Valentin et al., 2007) | VDO2007 | 19 | 6 | 5031 | 1 |
| Valentin, V. V., & O'Doherty, J. P. (2009) (Valentin & O'Doherty, 2009) | VO2009 | 17 | 1 | 5227 | 1 |
| Wulff, D. U., & Hertwig, R. | WH2012 | 186 | 6 | 15106 | 1 |
| Wulff, D. U. et al. (2015) (Wulff et al., 2015b) | WHH2015a | 63 | 10 | 10419 | 1 |
| Wulff, D. U. et al. (2015) (Wulff et al., 2015a) | WHH2015b | 124 | 32 | 49610 | 2 |
| Worthy, D. A., & Maddox, W. T. (2012) (Worthy & Maddox, 2012) | WM2012 | 114 | 2 | 9120 | 1 |
| Worthy, D. A., & Maddox, W. T. (2014) (Worthy & Maddox, 2014) | WM2014 | 66 | 3 | 16500 | 3 |
| Worthy, D. A. et al. (2008) (Worthy et al., 2008) | WMM2008 | 60 | 6 | 4740 | 2 |
| Worthy, D. A. et al. (2018) (Worthy et al., 2018) | WOCD2018 | 33 | 2 | 8250 | 1 |
| Wunderlich, K. et al. (2009) (Wunderlich et al., 2009) | WRO2009 | 23 | 1 | 6859 | 1 |
| Yechiam, E., & Busemeyer, J. R. (2006) (Yechiam & Busemeyer, 2006) | YB2006 | 80 | 2 | 32000 | 1 |
| Yechiam, E., & Busemeyer, J. R. (2008) (Yechiam & Busemeyer, 2008) | YB2008 | 180 | 6 | 107800 | 2 |

Continued on next page

| Article | Label | $N$ participants | $N$ problems | $N$ trials | $N$ studies |
|---|---|---|---|---|---|
| Yechiam, E. et al. (2005) (Yechiam et al., 2005) | YBE2005 | 24 | 1 | 2400 | 1 |
| Yechiam, E. et al. (2008) (Yechiam et al., 2008) | YDE2008 | 132 | 3 | 52800 | 2 |
| Yechiam, E., & Ert, E. (2007) (Yechiam & Ert, 2007) | YE2007 | 48 | 8 | 19200 | 2 |
| Yechiam, E., & Hochman, G. (2013) (Yechiam & Hochman, 2013) | YH2013 | 170 | 6 | 21700 | 2 |
| Yechiam, E. et al. (2015) (Yechiam et al., 2015) | YZA2015 | 188 | 8 | 78600 | 2 |
| Zaghloul, K. A. et al. (2012) (Zaghloul et al., 2012) | ZWLJ2012 | 12 | 2 | 2180 | 1 |
